# Supramolecular assembly of chloroplast NAD(P)H dehydrogenase-like complex with photosystem I from *Arabidopsis thaliana*

**DOI:** 10.1101/2021.12.16.472899

**Authors:** Xiaodong Su, Duanfang Cao, Xiaowei Pan, Lifang Shi, Zhenfeng Liu, Luca Dall’Osto, Roberto Bassi, Xinzheng Zhang, Mei Li

## Abstract

Cyclic electron transport/flow (CET/CEF) in chloroplasts is a regulatory mechanism crucial for optimization of plant photosynthetic efficiency. CET is catalyzed by a membrane-embedded NAD(P)H dehydrogenase-like (NDH) complex containing at least 29 protein subunits and associating with photosystem I (PSI) to form the NDH-PSI supercomplex. Here we report the 3.9 Å resolution structure of *Arabidopsis thaliana* NDH-PSI (AtNDH-PSI) supercomplex. We have constructed structural models for 26 AtNDH subunits, among which 11 subunits are unique to chloroplast and stabilize the core part of NDH complex. In the supercomplex, one NDH can bind up to two PSI-LHCI complexes at both sides of its membrane arm. Two minor LHCIs, Lhca5 and Lhca6, each present in one PSI-LHCI, interact with NDH and contribute to the supercomplex formation and stabilization. Our results showed structural details of the supercomplex assembly and provide molecular basis for further investigation of the regulatory mechanism of CEF in plants.

---

Photosynthetic electron transport is driven by light energy and can be classified into two types, the linear electron transport/flow (LET/LEF) and the cyclic electron transport/flow (CET/CEF)^1,2^. The LEF pathway involves both photosystems I and II (PSI and PSII) as well as cytochrome *b*_*6*_*f* (Cyt*b*_*6*_*f*), and produces ATP and NADPH, which are utilized in dark metabolic reactions, chiefly nitrate reduction and the Calvin-Benson cycle for CO_2_ assimilation^3^. As for the CEF pathway, electrons are recycled between Cyt*b*_*6*_*f* and PSI, generating the proton gradient across the thylakoid membrane^4,5^, which drives the ATP synthesis, without producing NADPH. CEF is crucial in balancing the ATP/NADPH ratio, since the ratio produced through LEF alone does not meet the requirement of the CO_2_ assimilation. Besides, phototrophs increase their ATP demands under stress conditions^6–8^. Moreover, the increased transmembrane delta pH produced by CEF is pivotal for inducing the energy-dependent quenching, which protects plants from photodamage^7,9,10^. Hence CEF plays essential roles in optimizing the growth and fitness of phototrophs^11,12^. In cyanobacteria and plants, type-I NAD(P)H dehydrogenase-like (NDH) complex (also called photosynthetic complex I) is crucial for CEF pathway, mediating the electron transport between PSI and Cyt*b*_*6*_*f*^13,14^. Previous genome analysis on NDH from cyanobacteria and plants^15,16^ indicated that 11 *ndh* genes encode homologous subunits (NdhA-K) of respiratory complex I^17^. However, NDH complex is different from respiratory complex I as it binds ferredoxin (Fd) as the electron donor instead of NADH^18–20^. In cyanobacteria, the large NDH complex (NDH-1L) possesses eight oxygenic photosynthesis-specific (OPS) subunits^21^, namely NdhL-NdhQ, NdhS and NdhV, in addition to the 11 conserved proteins^4^. Except for NdhQ, the other 18 subunits in cyanobacterial NDH-1L are conserved in plant NDH complex, which further contains more than 10 subunits specific to chloroplasts^4,22^. The cryo-electron microscopy (cryo-EM) structures of NDH-1L from *Thermosynechococcus elongatus* (TeNDH-1L) showed an L-shaped fold composed of a membrane and hydrophilic arms^18,19,23,24^, whereas plant NDH complex, which is larger and more complex, has not been structurally characterized at high resolution.

In angiosperms, the NDH complex is composed of at least 29 subunits, which can be divided into five subcomplexes^22,25^. The subcomplexes M and A (SubM and SubA) contain seven (NdhA-NdhG) and eight (NdhH-NdhO) subunits, corresponding to the membrane and hydrophilic arm of cyanobacterial NDH-1L, respectively. Subcomplex E (SubE) is composed of four subunits (NdhS-NdhV) and considered to be essential for binding and oxidizing Fd. Indeed, cyanobacteria have two SubE homologs, NdhV and NdhS, which were previously shown to be involved in Fd binding^18,19^. Subcomplex B (SubB) contains five subunits (PnsB1-PnsB5), which are conserved in chloroplast NDH and were suggested to form the second hydrophilic arm. The luminal subcomplex (SubL) is constituted of five subunits (PnsL1-PnsL5), which are suggested to be unique in the angiosperms^25,26^. Most of these chloroplast-specific subunits are crucial for the stability and/or activity of chloroplastic NDH.

In addition, photosynthetic NDH is able to associate with PSI to form NDH-PSI supercomplex^27,28^. In plants, the supercomplex formation increases the stability of chloroplastic NDH, especially under high light conditions, and therefore is critical for facilitating the CEF process *in vivo*^26,29,30^. Moreover, NDH-mediated CEF around PSI was shown to be the major contributor to PSI photoprotection in the presence of far red (FR) light^31^. In angiosperms, NDH is able to bind multiple copies of PSI^32^, and while the most populated form, the NDH–PSI_2_ supercomplex, is composed of one NDH sandwiched by two copies of PSI (referred as left and right PSI when viewed from the stromal side and placing SubA at the top) as revealed by the previous negative staining projection maps^32,33^. The two PSIs associate with NDH through their light-harvesting complex I (LHCI) belt, which usually constitutes Lhca1-a4 and Lhca2-a3 heterodimers^34,35^. However, the formation of NDH-PSI_2_ supercomplex largely depends on two minor LHCIs, Lhca5 and Lhca6^36^. They were suggested to locate in between the left PSI and NDH, or each replaces one major Lhca in the LHCI belt of each PSI, mediating the interactions with NDH^29,33^. The previously reported biochemical data and low-resolution models have provided important basis for understanding the subunit arrangement of NDH and its interaction with PSI, and yet, the detailed structure of the NDH-PSI supercomplexes is still lacking, which limits further exploration of the supercomplexes and understanding of the mechanism of CEF. Here we report the cryo-EM structure of *Arabidopsis* NDH-PSI_2_ supercomplex and reveal detailed information on the structure, location and assembly of subunits within this supercomplex, shedding light on the supramolecular basis of CEF process in plants.

## Overall structure

We purified NDH-PSI supercomplexes (Fig. S1) from *Arabidopsis* mutant strain NoM^37^ or from maize. Our cryo-EM analysis indicated that both smaller and larger particles, corresponding to NDH-PSI_1_ and NDH-PSI_2_, respectively, exist in maize (Fig. S2), while in *Arabidopsis* NoM strain, only NDH-PSI_2_ supercomplex was observed (Fig. S3). In agreement with previous result of the supercomplex from barley^33^, the only copy of PSI in the smaller complex is exclusively located at the left side of NDH, suggesting a more stable binding of the left PSI with NDH than the right copy. The three reconstruction maps show almost identical features, hence we used AtNDH-PSI supercomplex for further structural analysis, since the *Arabidopsis* mutant strain contains higher ratio of NDH-PSI_2_ particles (Figs. S2,S3). In addition, most of the biochemical data and mutant strains on CEF were obtained from *Arabidopsis thaliana*, a widely-studied model organism of plants. We finally solved the cryo-EM structure of AtNDH-PSI_2_ (hereafter referred to NDH-PSI) at an overall resolution of 3.9 Å (Fig. S3-S4, Table S1). Further focused refinement^38^ on two PSIs and two NDH regions (SubM-SubB-SubL and SubA-SubE) improved the resolution of these individual parts to 3.3 Å, 3.3 Å, 3.6 Å and 4.4 Å, respectively (Fig. S3). The structure shows that one NDH complex is sandwiched by two PSIs (Fig. 1a,b), consistent with early negative staining results^32,33^. No densities corresponding to additional Lhca proteins were observed. A total of 58 protein subunits and 423 cofactors were identified in the supercomplex structure (Table S2).

**Figure 1.**
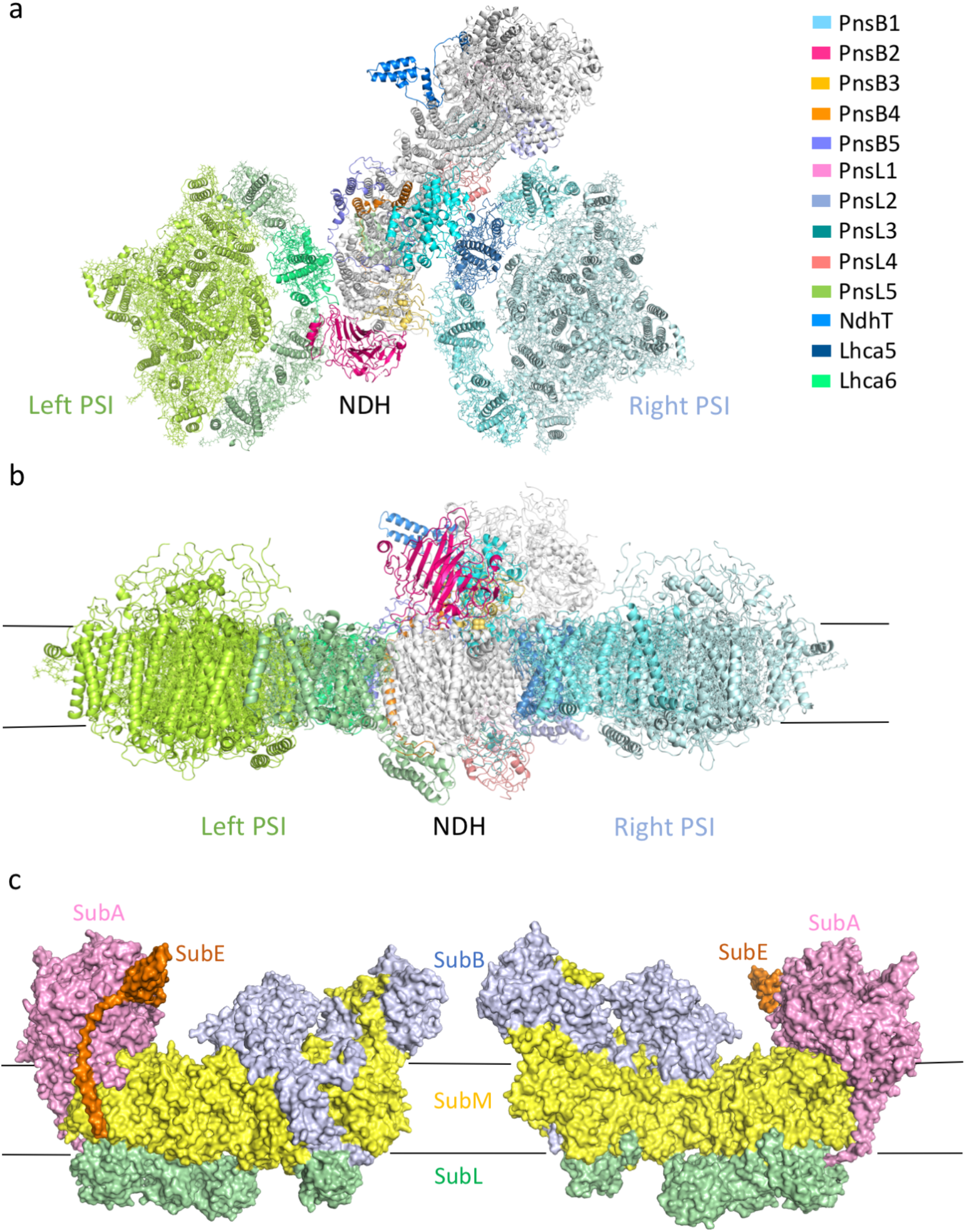
Overall structure of AtNDH-PSI supercomplex. (a) Cartoon representation of NDH-PSI supercomplex viewed from the stromal side. PnsB1-PnsB5, PnsL1-PnsL5 and NdhT (tentatively assigned) are colored differently, while subunits in SubM and SubA are colored white. Lhca6 and Lhca5 are highlighted in green and blue, in the left and right PSI, respectively. (b) Side view of NDH-PSI supercomplex. Color code is the same as in (a). (c) Surface representation of AtNDH complex. Five Sub complexes are colored differently and indicated.

In the NDH moiety, we modeled 26 out of the 29 subunits (Fig. 1c, Fig, 2, Fig. S5a), while three SubE subunits were not observed. The architecture of the core part of AtNDH (SubM and SubA) is similar with that of TeNDH-1L, forming an L-shaped structure (Fig. S5a). Five soluble subunits from SubL (PnsL1-PnsL5) cover the luminal surface of SubM, and five SubB subunits (PnsB1-PnsB5) are located at the distal region of NDH complex (Fig. 2). Additional density was observed close to SubA, and was tentatively assigned as one SubE subunit NdhT, which was built as poly-Ala chain (Fig. S4).

**Figure 2.**
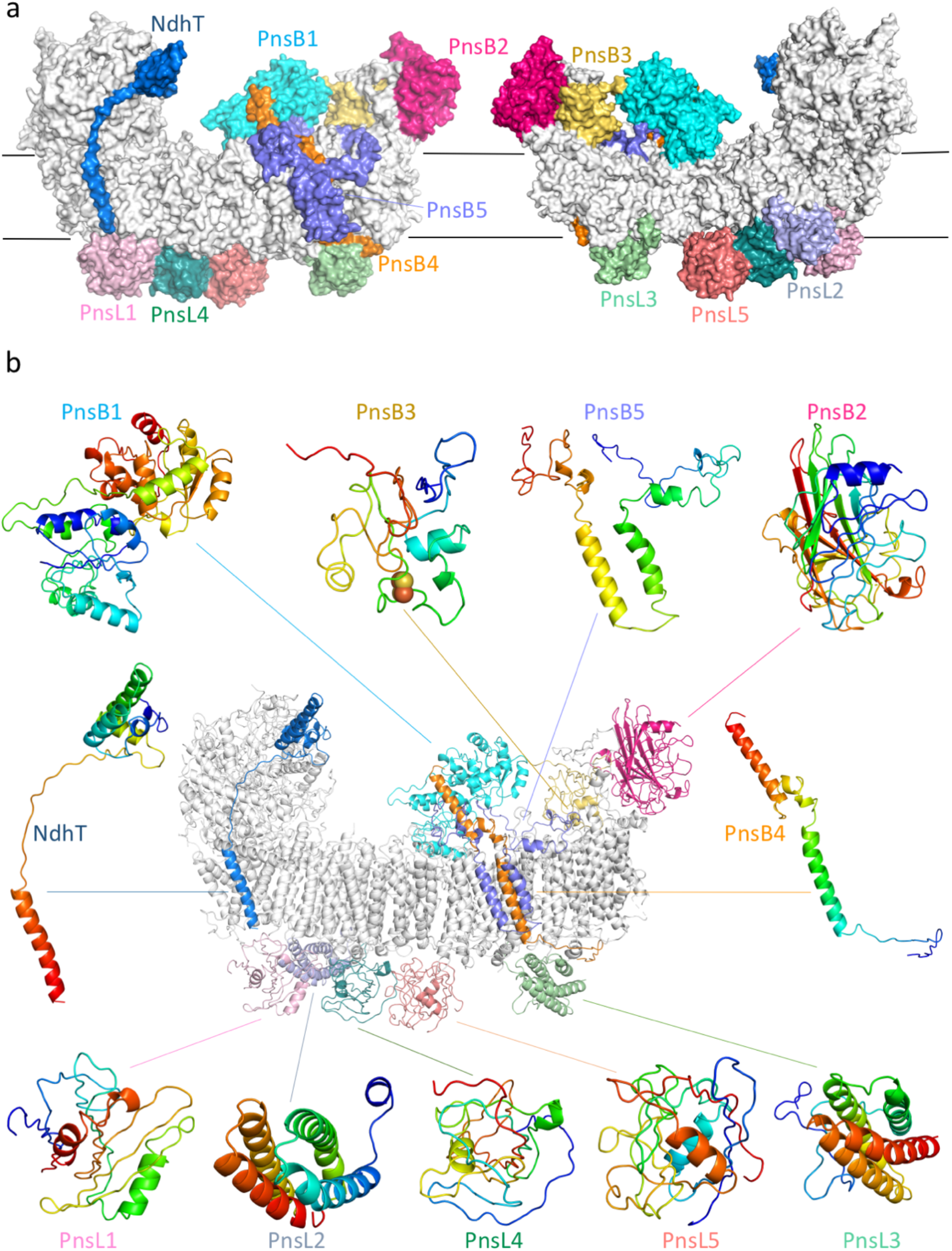
Structure and location of chloroplast-specific subunits in AtNDH complex. (a) Surface representation of NDH complex. PnsB1-PnsB5, PnsL1-PnsL5 and NdhT (tentatively assigned) from SubE are colored differently and labeled, while other NDH subunits are colored white. (b) Structures of PnsB1-PnsB5 from SubB, PnsL1-PnsL5 from SubL and NdhT. Structures of each single subunit are colored in rainbow mode, with blue and red indicating the N- and C-terminus.

The two PSI complexes adopt overall structures highly similar to the free PSI reported previously^39,40^. Each PSI contains 12 core subunits (PsaA-PsaL) and four LHCIs, which face NDH (Fig. 1a). The high-quality cryo-EM densities (Fig. S4) allow the accurate assignment of each subunits. Our structure shows that Lhca6 substitutes Lhca2 in the left PSI and contacts NdhF, whereas Lhca5 replaces Lhca4 in the right PSI and is at the vicinity of NdhD, consistent with previous functional studies^32^,

### SubM of AtNDH

In AtNDH complex, seven subunits (NdhA-NdhG) constitute the membrane arm core. While the overall architecture of SubM is similar to that of TeNDH-1L, AtNDH is more curved towards the right, which may allow a better match of the concave of SubM and the convex of LHC belt from the right PSI (Fig. S5b). Such a shape adaptation may facilitate the association of NDH with the right PSI complex.

The loop regions of SubM subunits were adapted to associate with chloroplast-specific NDH subunits. The NdhF exhibits the most evident differences between AtNDH and TeNDH-1L structures. AtNdhF possesses two insertions (loop-1 and loop-2) at the stromal region, which are crucial for stabilizing SubB (Fig. 3a,b, Fig. S6). Loop-1 forms an L-shaped coiled structure, protruding into the stromal space and providing the docking site for PnsB2 and PnsB3. Loop-2 extends outward from the distal end, forming a short amphiphilic helix, which is nearly parallel with the membrane plane, and forms close contacts with PnsB2 (Fig. 3b). Following this helix, a long horizontal helix (H-helix) spans NdhF, NdhD and NdhB, stabilizing the membrane arm of the complex. Curiously, although H-helix exhibits clear density, the C-terminal transmembrane helix (TMH16) of AtNdhF, which immediately follows the H-helix, does not show its density in the map (Fig. S7a). In contrast, in TeNDH-1L structure, the TMH16 is well resolved and clustered with NdhB, NdhG and NdhE (Fig. S7b), serving to stabilize these subunits and anchor the H-helix. In AtNDH, however, subunits from SubB and SubL interact with SubM at both the stromal and luminal side; thus, the TMH16 is likely not required for NDH-PSI supercomplex stabilization. Moreover, the large curvature of the membrane arm of AtNDH leaves little space to accommodate TMH16 (Fig. S7b). These structural features may result in the high flexibility of TMH16 in AtNDH.

**Figure 3.**
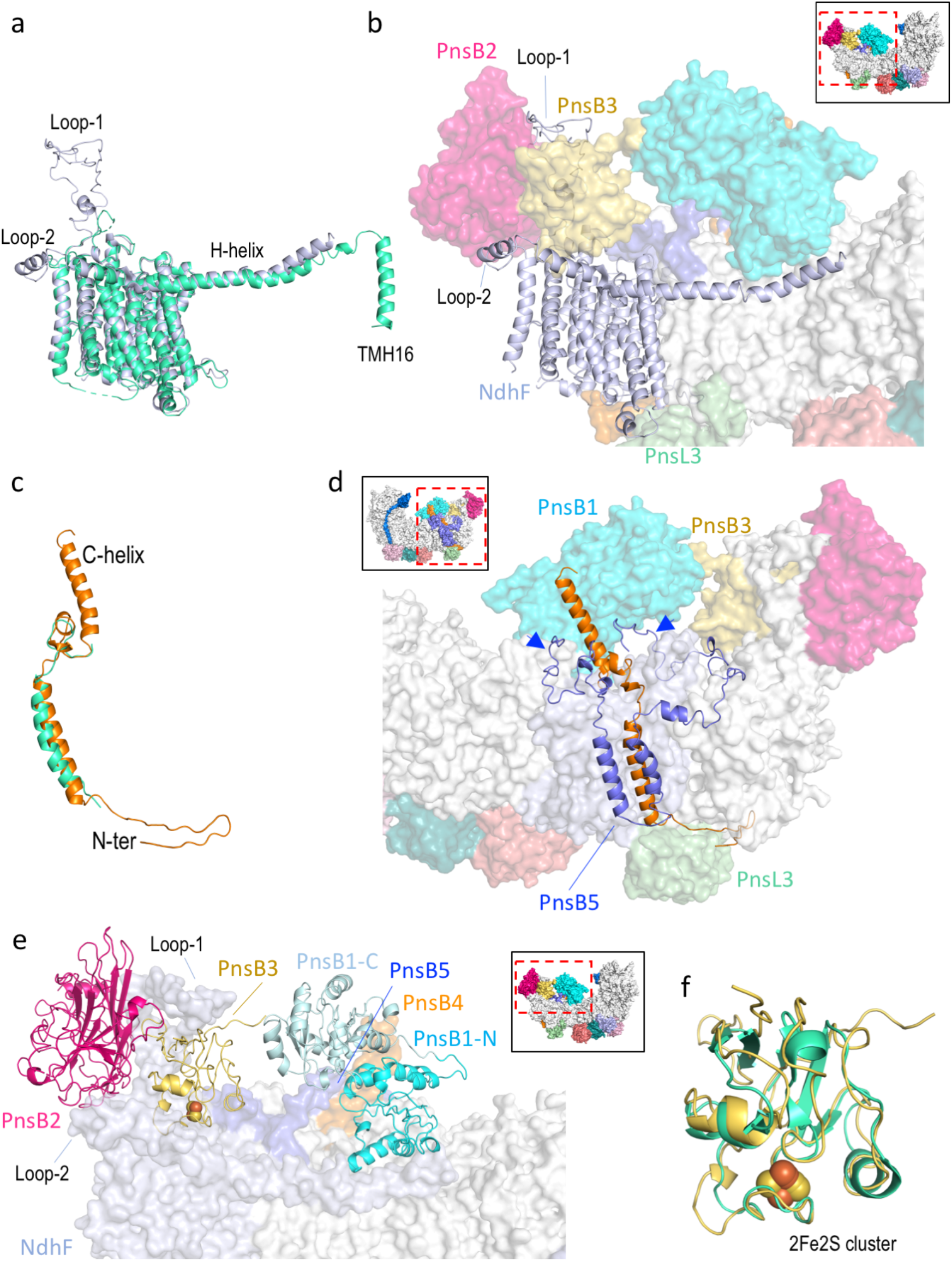
Interactions between SubB and SubM. (a) Structural comparison of AtNdhF (blue-white) and TeNdhF (green-cyan). Two insertions of AtNdhF (loop-1 and loop-2) as well as the H-helix and TMH16 (in TeNdhF) are indicated. (b) Interaction between NdhF and SubB. NdhF is shown in cartoon mode and in the same view as in (a), while other subunits are in surface mode. Two insertions of AtNdhF (loop-1 and loop-2) provide binding sites for PnsB2 (magenta) and PnsB3 (yellow). (c) Structural comparison of AtPnsB4 (orange) and TeNdhP (green-cyan). The C-helix and the N-terminus of PnsB4 are indicated. (d) Interaction of PnsB4 (orange cartoon) and PnsB5 (slate cartoon) with other subunits (surface mode). The transmembrane helices of PnsB4 and PnsB5 attach to NdhD (blue-white). PnsB4 interacts with PnsB1 (cyan) via its C-helix, and with PnsL3 (light-green) through its N-terminal tail. PnsB5 has two elongated terminal loops (indicated by blue arrows), encircling the NdhD subunit (blue-white) at the stromal side, and the N-terminal loop simultaneously contacts PnsB1 (cyan). (e) Interaction of PnsB1, PnsB2 and PnsB3 (cartoon mode) with other subunits (surface mode). PnsB2 (magenta) and PnsB3 (yellow) cluster together and are supported by Loop-1 of NdhF (blue-white). PnsB1-N (cyan) attaches to the membrane arm, while PnsB1-C (pale-cyan) away from the membrane and interacts with PnsB3 (yellow). (f) Structural comparison of PnsB3 (yellow) and Fd (PDB code 1L5P) (green-cyan). The 2Fe2S cluster in two structures are superposed well and labeled.

### SubB of AtNDH

The five SubB subunits are essential for NDH activity, lacking one leads to the defect in accumulation of the NDH complex^36,41–43^. Among them, three (PnsB1-PnsB3) are soluble proteins located at the stromal side, and the remaining two (PnsB4-PnsB5) are membrane-embedded subunits. In the structure, SubB can be divided into two parts, namely PnsB1-4-5 associated with NdhD and PnsB2-3 (located at the distal end of SubM) attached on the top of NdhF (Fig. 2).

PnsB4 is the only SubB subunit that has a counterpart (NdhP) in cyanobacterial NDH-1L. Both PnsB4 and NdhP have one TMH, and attach to the periphery of NdhD. However, PnsB4 is considerably larger than NdhP, with extensions in the N- and C-terminal regions (Fig. S8a). The N-terminal extension forms a hairpin loop, which is almost parallel with the luminal β-sheet (β-H) of NdhF, and simultaneously contacts PnsL3. The C-terminal extension forms a long α-helix (C-helix) protruding into the stromal side and interacting with PnsB1 (Fig. 3c,d). PnsB5 contains two TMHs flanking at the periphery of PnsB4. Two elongated arm-like N/C-terminal regions of PnsB5 reach out to embrace the stromal surface of NdhD (Fig. 3d).

At the stromal side, PnsB1 is the largest SubB subunit of AtNDH and can be divided into N- and C-terminal domains (PnsB1-N and PnsB1-C) (Fig. 3e). The PnsB1-N strongly interacts with NdhD, NdhB and H-helix of NdhF at the stromal side. Moreover, the C-helix of PnsB4 and N-terminal tail of PnsB5 attach to the interface between PnsB1-N and PnsB1-C, and the latter further contacts PnsB3 (Fig. 3e). PnsB3 and PnsB2 are clustered together and clamped by the two insertions of NdhF (Fig. 3e). PnsB2 is mainly composed of β-sheets and binds at the tip of the membrane arm (Fig. 2). We also solved the crystal structure of PnsB2 protein at 2.55 Å resolution (Fig. S8b). Structural comparison showed that a loop region (aa. 263-272) in PnsB2 adopts different conformation in the complex which facilitates its interaction with NdhF and PnsB3 (Fig. S8b). Hence the five SubB subunits connect with each other, and form extensive interactions with SubM both within and outside the thylakoid membrane. The strong association within SubB and between SubB and SubM/PnsL3 may explain the crucial role of SubB subunits in NDH accumulation and function^36,41–43^.

Furthermore, PnsB3 was previously suggested to host an iron-sulfur cluster ^41^, and we did observe the density corresponding to a 2Fe-2S cluster (Fig. S4), which is located right above the membrane surface plane (Fig. 3e) and faces the right PSI. Surprisingly, although PnsB3 does not share close sequence similarity with ferredoxin (Fig. S8c), it adopts a ferredoxin-like structure, with the 2Fe-2S cluster positioned at a pocket similar to the one in ferredoxin (PDB code 1L5P) (Fig. 3f). The iron-sulfur cluster in PnsB3 was previously shown to be redox-active, suggesting a possibility that PnsB3 might be able to sense the redox state in the stromal region, or even participates in the electron transport process.

### SubL of AtNDH

All five SubL subunits are soluble proteins and line up at the luminal surface of SubM (Fig. 1c). PnsL1 and PnsL2/PnsL3 resemble two oxygen-evolved complex (OEC) subunits of plant PSII, PsbP and PsbQ (Fig. S9a,b), respectively^44–46^. PnsL4 and PnsL5 are two immunophilins, belonging to FKBP and cyclophilin subfamilies, respectively^36,47^. The OEC subunits have crucial roles in stabilizing the functional conformation of PSII^48^. Immunophilins usually show peptidyl-prolyl cis-trans isomerase (PPIase) activity^49,50^ and serve as scaffolding proteins for supercomplex assembly. These results suggest that the SubL assists the functional assembly and organization of chloroplastic NDH complex. In agreement with the suggestion, previous biochemical and genetic studies demonstrated that the SubL subunits, except PnsL5^47^, are crucial for the accumulation and activity of the chloroplast NDH complex in thylakoids^36,44–46^.

In the structure, PnsL3 is separated from other SubL subunits and joins SubB to stabilize NdhD and NdhF. It interacts with the N-terminal tail of PnsB4 in addition to NdhF and NdhD (Fig. 4a,b), in agreement with the earlier observation that PnsL3 requires PnsB4 in order to associate with NDH complex^44^. Other SubL subunits are clustering together. PnsL1 and PnsL2 strongly associate with each other and attach to the luminal face of NdhA-C-E-G. PnsL4 is located beneath NdhB and interacts with PnsL1 and PnsL2 simultaneously. Together, the three SubL subunits (PnsL1, PnsL2 and PnsL4) form a triangular structure (Fig. 4a). PnsL5 strongly associates with PnsL4 and interacts with NdhD via its N-terminal fragment (Fig. 4b). Therefore, SubL interacts closely with the SubM subunits (Fig. 1c), and together with SubB, they may facilitate the coupling of proton translocation processes performed by SubM subunits in AtNDH.

**Figure 4.**
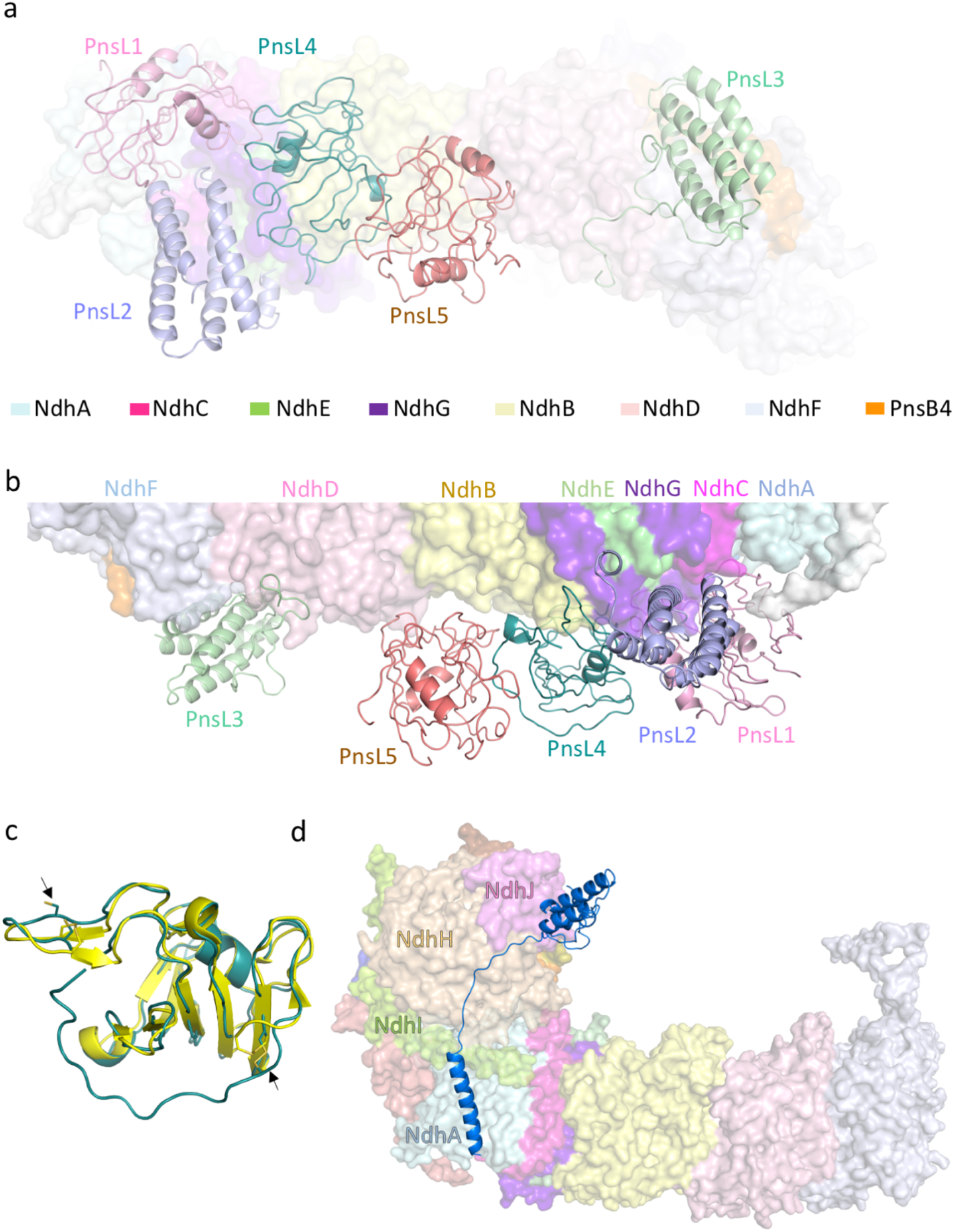
Interactions of SubL and NdhT with other subunits. (a) Luminal view of SubL subunits in AtNDH. SubL subunits are shown in cartoon mode, while other subunits are in surface mode. PnsL1 (pink), PnsL2 (light-blue) and PnsL4 (deep-teal) form a triangular structure, interacting with NdhA-C-E-G-B. PnsL5 (deep-salmon) interacts with PnsL4 and covers the luminal surface of NdhB. PnsL3 (light-green) is separated from other SubL subunits, and close to PnsB4 (orange), NdhF (light-blue) and NdhD (light-pink). (b) Sideview of SubL (cartoon mode) and SubM (surface mode) in AtNDH. A void space is present between PnsL5 and SubM. (c) Comparison of PnsL4 (deep-teal) and AtFKBP-13 (PDB code 1U79) (yellow). The conserved cysteine residues are shown in sticks and indicated by black arrows. (d) Side view of NdhT (cartoon mode) and the core part of AtNDH (SubM and SubA, surface mode). NdhT interacts with Ndh-H-I-A.

In the structure, PnsL5 covers the luminal surface of NdhB; yet does not seem to have any direct contacts: a large void space is present between them (Fig. 4b). Their loose association is consistent with previous result that lack of PnsL5 does not result in severe dysfunction of NDH^47^. Interestingly, PnsL4 shows high sequence and structural similarity with another FKBP family member AtFKBP13 (1U79) (Fig. 4c, Fig. S9c), which was previously suggested to interact with the Rieske subunit of Cyt*b*_*6*_*f* through its N-terminal signal peptide^51,52^. PnsL4 possesses a longer N-terminal tail than AtFKBP13 (Fig. S9c), which might be able to bind Rieske subunit, potentially allowing NDH to form direct interaction with Cyt*b*_*6*_*f*. Additionally, PnsL4 possesses two pairs of cysteine residues potentially form disulfide bonds at the same positions as AtFKBP13 (Fig. S9c). Hence PnsL4 may sense the redox signal in the lumen and respond to the redox change so as to regulate the assembly and/or functional activity of NDH.

### SubA and SubE of AtNDH

SubA contains eight NDH subunits (NdhH-NdhO), among which, NdhL is the only membrane subunit, but attributed to SubA due to its need for the stabilization of SubA in AtNDH^53^. All SubA subunits are highly similar with their counterparts in TeNDH-1L, in both sequence and overall fold. In the structure, the SubA exhibits lower resolution of ~4.4 Å (Fig. S3), which may reflect its higher mobility.

SubE attaches to SubA and is the most poorly defined region in our structure. SubE is composed of four subunits (NdhS-NdhV). Among them, NdhS and NdhV have counterparts in cyanobacterial NDH-1L, but were not observed in our structure. The other two are chloroplast-specific subunits NdhT and NdhU. Both were suggested to possess a J(-like) domain similar to molecular chaperone DnaJ and a C-terminal TMH^20^. Previous data suggested that NdhT associates with SubA and is required for the accumulation of SubA, whereas NdhU is at the periphery of NdhT^20^. The cryo-EM density corresponding to a J-domain followed by a long loop and a TMH was observed at the periphery of NDH, contacting SubA at the side of NdhJ, NdhH, NdhI and NdhA (Fig. 4d, Fig. S4). This subunit is probably attributed to NdhT or NdhU. We tentatively assign it as NdhT, based on the previous suggestion that NdhU is located on the outside of NdhT^20^, and on our finding that the secondary structure of NdhT predicted by AlphaFold^54^ fits the density better. Interestingly, mitochondrial complex I also possesses a supernumerary subunit, containing a single TMH, which is located at the same position of NdhT (Fig. S10), and strictly required for assembly of a functional complex I ^55^. These structural observations suggest that NdhT may have an important role in NDH assembly and activity^20^.

### NDH-PSI Assembly

In the supercomplex, the left PSI contains Lhca1-a4 and Lhca6-a3 dimers, with Lhca6 substituting Lhca2 found in the free PSI. Lhca6 shows similar structure as Lhca2, but it adopts different conformation in the N-terminal loop and the loop connecting the second and third TMHs (AC-loop) (Fig. 5a, Fig. S11a). The different conformation of the AC-loop results in loss of one chlorophyll (Chl 618) in Lhca6 when compared to Lhca2 and weaker association between Lhca1-a4 and Lhca6-a3 heterodimers (Fig. 5a,b). In Lhca6, both the AC-loop and N-terminal fragment are extending to the stromal region and towards NDH complex, thus are pivotal for the NDH-Lhca6 interaction. Part of the AC-loop inserts into the interface between NdhF and PnsB2, while other region of AC-loop and the elongated N-terminal tail of Lhca6 interact with NdhF and PnsB5, respectively (Fig. 5c). This structural feature is consistent with previous suggestion that SubB is the contacting site for Lhca6^29,36^. Moreover, the luminal region of Lhca6 and the N-terminus of NdhF are close to each other (Fig. 5d) and form Van der Waal contacts. The strong interaction between Lhca6 and NDH may explain the previous finding that Lhca6 was not detected in free PSI but only localized in NDH-PSI supercomplex^36,56^.

**Figure 5.**
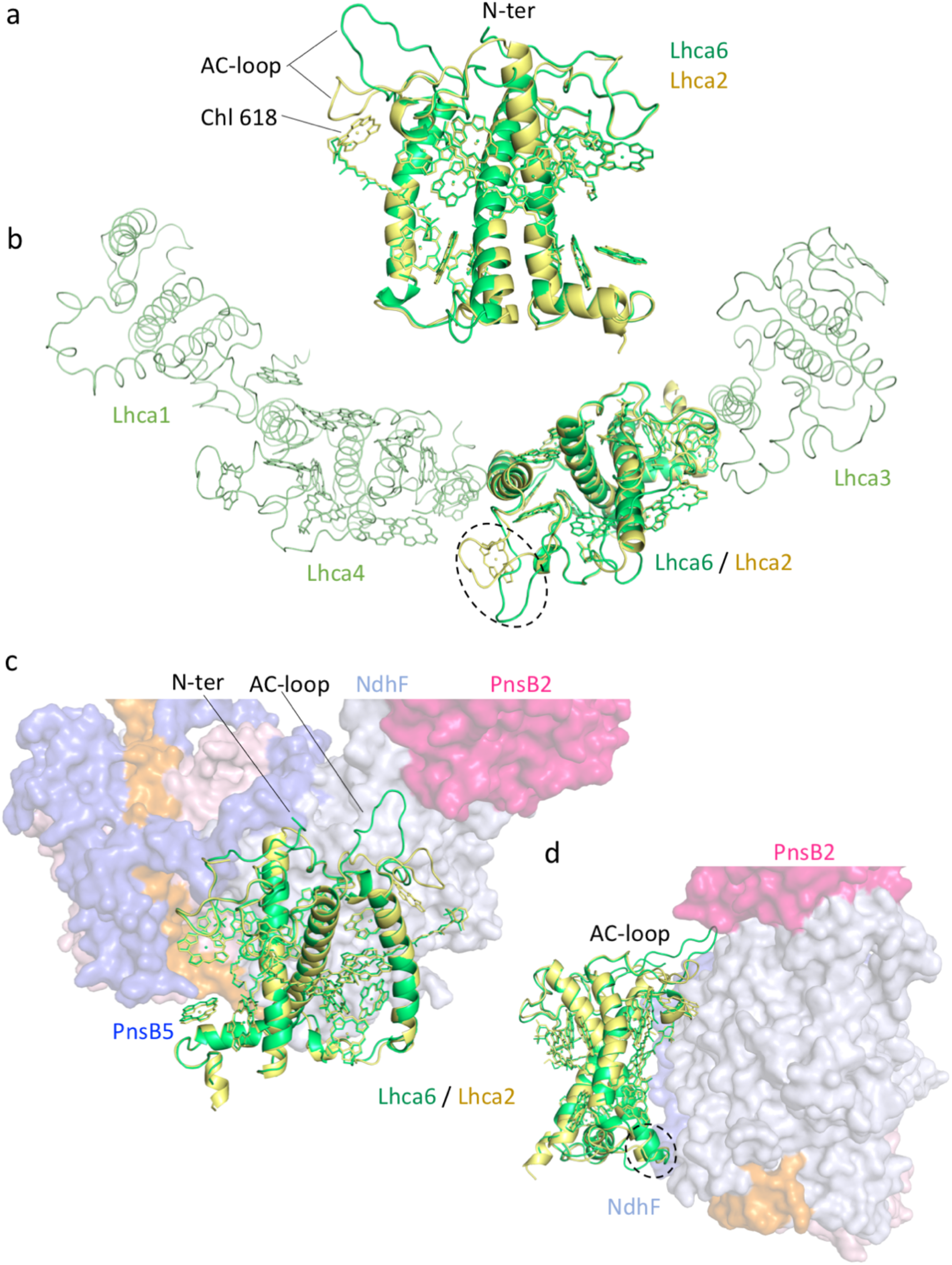
The left PSI and NDH. (a) Structural comparison of AtLhca6 and Lhca2 from maize PSI (ZmLhca2) (PDB code 5ZJI). Pigment molecules are shown in stick mode, and the phytol chains of chlorophylls are omitted for clarity. The AC-loop and N-terminal tail of Lhca6 adopt different conformation, and Lhca6 lacks Chl 618 comparing with Lhca2. (b) Stromal side view of the LHCI belt. ZmLhca2 was superposed on AtLhca6 for comparison. The differences around the AC-loop of AtLhca6 (indicated by dashed circle) may result in the weaker interaction between Lhca6-a3 and Lhca1-a4 dimers. (c,d) Side view of NDH (surface mode) and Lhca6 (cartoon mode) from two different angles. ZmLhca2 was superposed on AtLhca6 for comparison. Both AC-loop and N-terminal tail of AtLhca6 are involved in the interaction with NDH at the stromal side (c), and the luminal regions of AtLhca6 is close to NdhF (indicated by dashed circle), and form hydrophobic interactions (d).

The right PSI contains Lhca1-a5 and Lhca2-a3 dimers, with Lhca5 replacing Lhca4 found in the free PSI. Several hydrophobic interactions were observed at the stromal surface between PnsB1 and the AC-loop of Lhca5, and Van der Waal contacts were found between NdhF and Lhca2. Lhca5 has two insertions in its AC-loop compared with Lhca4 (Fig. S11b), which fold backwards to avoid clash with PnsB1, thus can better associate with PnsB1 and Lhca1 (Fig. 6a,b). The right PSI is loosely associated with NDH, as there is a large gap present between them (Fig. 6c). Lipid molecules may fill in the gap and mediate their interactions. Lhca5 does not form as strong interactions with NDH subunits as Lhca6, and may be able to bind within the LHCI belt in the absence of NDH. However, compared with Lhca4, Lhca5 forms a weaker dimer with Lhca1. Lhca5 lacks Chl 618 due to the conformational change of its AC-loop, and it does not contain Chl 617 owing to lack of a conserved ligand residue (Fig. S11b). Both Chl 617 and 618 are located at the Lhca1-a5 interface. Moreover, a lutein molecule, which associates with Chl 617 of Lhca4, located at the Lhca1-a4 interface, is lost in Lhca1-a5 dimer (Fig. 6b). The fewer chlorophylls in Lhca5 and weaker Lhca1-a5 dimer interaction (compared with Lhca1-a4) observed in our structure is consistent with previous results indicating that Lhca5 is present in substoichiometric amounts in free PSI ^57^.

**Figure 6.**
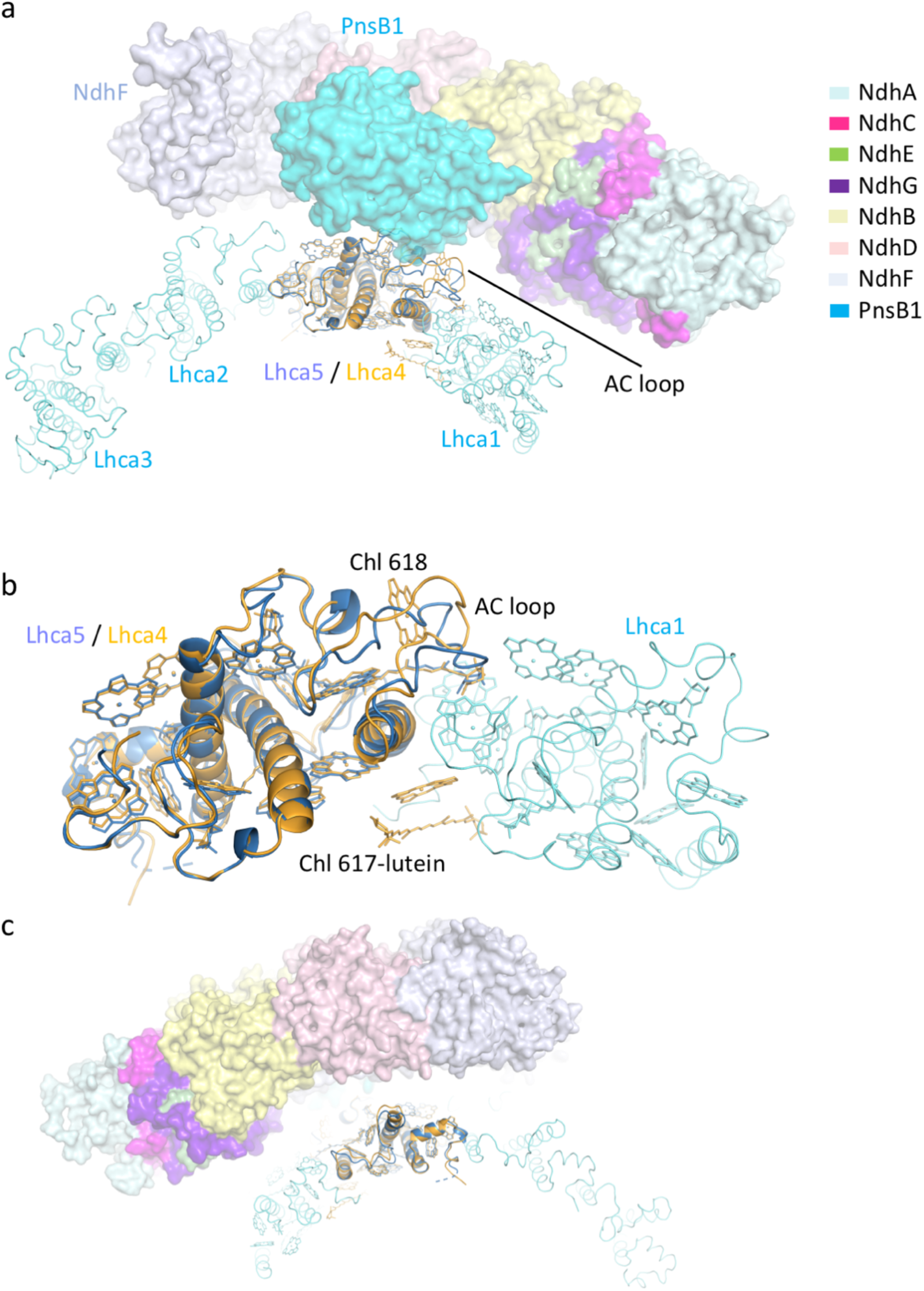
The right PSI and NDH. (a) Stromal side view of NDH (surface mode) and the LHCI belt of right PSI. ZmLhca4 (PDB code 5ZJI) was superposed on AtLhca5 for comparison and both are shown in cartoon mode. Other LHCIs are shown in ribbon mode. Pigment molecules are shown in stick mode, and the phytol chains of chlorophylls are omitted for clarity. The pigments in Lhca2-a3 dimer are also omitted for clarity. The AC-loop is indicated. (b) Stromal side view of the Lhca1-a5 dimer. ZmLhca4 was superposed on AtLhca5 for comparison. AtLhca5 lacks two chlorophylls (Chls 617 and 618) and one lutein molecule compared with ZmLhca4. (c) Luminal side view of NDH (surface mode) and the right PSI (cartoon and ribbon mode). A large space is present between NDH and the right PSI.

The structure showed that the association of both left and right PSI with NDH requires SubM and SubB, in agreement with previous results showing that without SubA, other subunits are still able to assemble with two copies of PSI^28,36^. Moreover, it was suggested that incorporation of SubA is the last step of NDH-PSI assembly process^58^, which may result in the relatively loose association of SubA with other parts of NDH, and explain why SubA exhibits weak density in our structure. The two PSIs are not equivalent when they associate with NDH, with the left PSI more strongly attached to NDH through Lhca6 (Figs. 5,6). This structural feature is consistent with earlier finding that Lhca6 is co-expressed with NDH subunits^36^, and explaining our observation on the cryo-EM results of maize NDH-PSI as well as the previous observation that the NDH–PSI_1_ supercomplex exists only in one form, in which the left PSI is preserved, whereas the right one is absent^33^.

### Functional implications

During CEF, PSI delivers electrons to Fd, which further transfers electrons to NDH and reduces the plastoquinone molecule bound within NDH complex. The plastoquinol diffuses to Cyt*b*_*6*_*f* and the electron is used to reduce the lumen-located electron-carrier plastocyanin (Pc). Subsequently, the reduced Pc leaves Cyt*b*_*6*_*f* and binds to PSI to deliver the electron. It was suggested that the formation of NDH-PSI supercomplex may enhance the CEF process by allowing a more efficient channeling of electrons^14^. Therefore, we have measured the binding affinity between NDH-PSI complex and fFd, and found that it falls into the similar range of the binding affinity between free PSI and Fd (Fig. S12). The strong interaction between PSI and Fd, and the short distance between PSI and NDH suggest that Fd could efficiently transfer electrons from PSI to NDH within the supercomplex. *Arabidopsis* NoM mutant has low PSII efficiency due to the disrupted connection between the PSII core and peripheral antennae^37^, which may affect the LEF process, hence requires a higher CEF activity. This might be the reason that most of the NDH complex binds two PSIs in this mutant. Our structure showed that the NDH-associated PSI contains fewer chlorophylls than the free PSI, due to the loss of chlorophylls in Lhca5 and Lhca6, suggesting that, during evolution, plants may sacrifice their light-harvesting capability to form NDH-PSI supercomplex for better CEF performance.

## Materials and Methods

### Sample purification and characterization for cryo-EM

NoM mutant of *Arabidopsis thaliana* (knocked out *lhcb4* and *lhcb5* gene)^37^ were grown about two months (50 µmol photons mx^−2^ s^−1^, 12 h photoperiod, 22°C) and chloroplast were isolated from its leaves as described previously^59^ with little modification. For purification of NDH-PSI supercomplex, the chloroplast was adjusted to chlorophyll concentration of 1.0 mg ml^−1^, then directly solubilized with equal volume of 1.8% dodecyl-β-D-maltoside (β-DDM) which add 0.5 mM Phenylmethanesulfonyl fluoride (PMSF), 0.5 mM Benzamidine and 2 mM ε-aminocaproic acid. After centrifugation at 20,000 g for 10 min, the supernatant was fractionated through sucrose-density gradient ultracentrifugation at 187,600 g, 4 °C for 18 h (Beckman SW41 rotor). The sugar solution contains 0.65 M sucrose, 25 mM Tricine (pH 7.8), 5 mM MgCl_2_, 10 mM KCl, 0.01% α-DDM, 0.02% digitonin. Maize were grown about two weeks (50 µmol photons m^−2^ s^−1^, 12 h photoperiod, 22°C), and the chloroplast isolation and NDH-PSI purification from maize are similar with that from *Arabidopsis*, except the sucrose gradient is 15%-45% (w/v), 25 mM Tricine (pH 7.8), 5 mM MgCl_2_, 10 mM KCl, 0.01% α-DDM, 0.02% digitonin and gradient ultracentrifugation at 246,000 g. The band containing NDH-PSI supercomplex were obtained (Fig. S1a) and concentrated to chlorophyll concentration of 1.5 mg ml^−1^ for cryo-EM specimen preparation.

The sodium dodecylsulfate polyacrylamide gel electrophoresis (SDS-PAGE) (Fig. S1b), western blotting (Fig. S1c), spectra measurements (Fig. S1d,e) and the high-performance liquid chromatography (HPLC) (Fig. S1f) analysis of the NDH-PSI samples used for cryo-EM were performed according to the previous report^60^.

### Sample preparation and data acquisition for cryo-EM

NDH-PSI sample was added to glow discharged holey carbon grid (GIG-C31213) and flash-frozen in liquid ethane at around 100 K using a semi-automatic plunge device (Thermo Fisher Scientific Vitrobot IV) with a blotting time of 3 s and blotting force of level 3 at 100% humidity, 4°C. All micrographs were collected using SerialEM data collection software^61^ with dose-fractionated super-resolution mode, with magnification of 22,500×, each exposure of 6.4 s was dose fractionated into 32 movie frames, leading to a total dose of ~60 e^−^ Å^−2^.

For the *Arabidopsis* NDH-PSI, a total of 7,148 movies were collected on a 300 kV FEI Titan Krios electron microscope (Thermo Fisher Scientific) equipped with K2 direct electron detector (Gatan), pixel size is 1.04. For maize NDH-PSI, a total of 3,438 movies were collected on a 300 kV FEI Titan Krios electron microscope equipped with K3 direct electron detector (Gatan), pixel size is 1.07.

### Cryo-EM data processing

All movie stacks were subjected to beam-induced motion correction using MotionCor2^62^. The parameters of the contrast transfer function were determined by the program CTFFIND4^63^. All subsequent processing steps were performed with RELION 3^64^. For the NDH-PSI complex from *Arabidopsis*, the reference based auto-picking generated a total number of 468,471 particles. After several rounds of 2D classifications, 337,219 particles were selected for further analysis of 3D classification using the map generated from the RELION initial model building as the reference model. 136,022 particles from two classes with good feature were kept for the following 3D auto-refinement, yielding a density map of the NDH-PSI complex with an overall resolution of 3.9 Å. To improve the density map, focus refinements were performed for the regions of PSI-1, PSI-2 and NDH-MBL, and reached a final resolution of 3.3 Å, 3.3 Å and 3.6 Å, respectively. For the region of NDH-AE, a no-alignment classification was applied after particle subtraction of the signal of other regions beyond the NDH-AE, and further focus refinement of the selected 76,085 particles resulted in a density map of 4.4 Å (Fig. S3). For the NDH-PSI complexes from Maize, the cryo-EM data processing flow was similar as above. Briefly, 3D auto-refinement of the 26,620 particles selected from the first round of 3D classification generated a density map of the complex of NDH bound to two PSI with a resolution of 4.5 Å. A subset of 10,343 particles from the second round of 3D classification represents the complex of NDH bound to one PSI, and further 3D auto-refinement resulted in a density map at 4.4 Å (Fig. S2). The resolutions were estimated based on the gold-standard Fourier shell correlation with 0.143 criterion. The local resolutions of the final density maps were measured using ResMap^65^ (Figs. S2,3).

### Cloning, expression, purification, crystallization and diffraction data collection of PsnB2

The full-length gene encoding PsnB2 of *Arabidopsis* were artificially synthesized (GenScript, China) and constructed to vector pMCSG9 following the manufacturer’s instructions. The plasmid contained 6*His, Maltose-binding protein (MBP) tags and TEV protease site before the PsnB2 gene in the N-terminus was transformed into *Escherichia coli* strain BL21 (DE3) (TransGen Biotech) and cultured on Luria-Bertani (LB) plate containing 100 µg ml^−1^ ampicillin. The clones were cultured in LB medium containing ampicillin (100 µg ml^−1^) at 37 °C and shaken at 220 rpm until the OD600 reached 1.0-1.3, then the protein expression was induced by adding 1 mM Isopropyl-β-D-thiogalactopyranoside (IPTG) overnight at 16 °C. The cells were collected at 6,000 rpm for 10 min, re-suspended in lysis buffer (25 mM Tris, pH 8.0, 300 mM NaCl, 5% glycerol, 1 mM TCEP) and disrupted by sonication, then the lysed cells were centrifuged at 18,000 rpm for 40 min. The supernatant was loaded onto Ni^2+^ affinity resin (Ni-NTA: GE healthcare) and washed with lysis buffer containing 30 mM imidazole. The target protein was eluted with 250 mM imidazole in lysis buffer. The protein was digested with TEV protease in 8:1 (w/w) ratio and dialysis with lysis buffer simultaneously at 4 °C overnight, then the mixture was loaded onto Ni-NTA affinity-chromatography again to remove the MBP and His-tag. The target proteins flowed through from the column were collected and further purified by size-exclusion chromatography (Superdex 200 10/300 GL, GE Healthcare) with lysis buffer containing 5 mM DTT. Then the proteins were concentrated to 10 mg/ml using the 30 kDa cut-off Amicon Ultra-4 centrifugal filter unit (Millipore), and stored at −80 °C for crystallization.

The PsnB2 crystallization trials were set up using a Hampton Research Screen Kit by applying the sitting drop vapor diffusion technique. The final optimized crystal was obtained by mixing equal volume of protein (8 mg ml^−1^ protein) and reservoir solution composed of 25% PEG3,350, 1 M Bis-Tris, pH6.5, 0.2 M Lithium sulfate. After incubation at 18 °C for two weeks, single crystals grew to a useful size and were flash-cooled in liquid nitrogen before diffraction data collection. The diffraction data were collected at BL17U of Shanghai Synchrotron Radiation Facility (SSRF)^66^ and processed by HKL2000 program^67^.

### Cloning, expression and purification of Ferredoxin(Fd)

*Arabidopsis* Fd gene were synthesized (GenScript, China) and constructed to PET28a, then transformed into *Escherichia coli* strain BL21 (DE3), after cultured on LB medium plate containing 50 µg ml^−1^ Kanamycin Monosulfate, the clones were cultured in LB medium (50 µg ml^−1^ Kanamycin Monosulfate) at 37 °C, 160 rpm to OD600 about 1.0, then add 1 mM IPTG and 1 mM FeSO_4_ for protein expression at 16 °C shaked about 20 h. The cells were harvested and re-suspended in lysis buffer (25 mM Tris, pH8.0, 300 mM NaCl, 5% glycerinum) and disrupted by sonication, after centrifuged, the supernatant was load onto Ni^2+^ affinity resin (Ni-NTA: GE healthcare), wash and elution buffer are 30 mM and 300 mM imidazole in lysis buffer. Before SPR assay, Fd was purified by size-exclusion chromatography (Superdex 200 10/300 GL, GE Healthcare) with 10 mM HEPES, 100 mM NaCl, then concentrated to 10 mg ml^−1^.

### Surface plasmon resonance (SPR)-based biosensor analysis

The banding affinity between *Arabidopsis* NDH-PSI and PSI with Fd was detected by SPR using a Biacore T100 biosensor (GE Healthcare). The experiment was performed as previously described with some modifications^19^. NDH-PSI and PSI was diluted with 10 mM NaAc (pH4.5) to the protein concentration of 50 µg mL^−1^, and then immobilized on the activated CM5 chip surface. The running buffer is 10 mM HEPES (pH7.5), 100 mM NaCl, 0.03% α-DDM. A series of concentration of Fd (10 µM, 5 µM, 2.5 µM, 1.25 µM, 0.625 µM, 0.3125 µM) were diluted using the running buffer, then injected using a flow rate of 30 µL min^−1^, contact time is 60 s, dissociation time is 120 s. Regeneration solution is 1 M NaCl and regenration time is 60 s. The data were analyzed with Biacore T100 evaluation software (GE Healthcare) by fitting to a 1:1 binding model (Fig S12). The experiment has been repeated three times with similar results.

### Model building and refinement

To construct the structural model of the NDH-PSI supercomplex from *Arabidopsis*, the PSI moieties from the maize PSI-LHCI-LHCII complex (PDB code: 5ZJI) was docked into the 3.9 Å map and the 3.3 Å focused refinement maps, respectively, using UCSF Chimera^68^. The amino acid residues of each subunits were mutated based on the sequence of *Arabidopsis* using Chainsaw in CCP4^69^. Lhca6 in PSI-1 and Lhca5 in PSI-2 were mutated manually according to the sequences and the density maps using Coot^70^.

To construct the structural model of NDH moiety, the model of the homologous subunits NdhA-NdhG from cyanobacteria NDH-1L structure (PDB code 6KHJ) was docked into the 3.6 Å focused refinement maps (NDH-MBL), and the model of NdhH-NdhO was docked into the 4.4 Å focused refinement maps (NDH-AE) using UCSF Chimera, respectively. The amino acid residues of each subunits were mutated based on the sequence of *Arabidopsis* using Chainsaw in CCP4^69^. Subunits PsnL1-PsaL5, PsnB1-PsnB5 are built manually based on the 3.6 Å focused refinement NDH-MBL map and the secondary structure prediction results by PSIPRED^71^. Because the densities of PsnB2 was not clear enough to trace the direction of βstrands precisely, we solved the crystal structure of PsnB2 at 2.55 Å by molecular replacement using the electron microscopy map as an initial model following the method reported in reference^72^. Then the crystal structure of PsnB2 was docked into the 3.6 Å focused refinement maps (NDH-AE) using UCSF Chimera^68^ for further refinement. For the model building of NdhT in NDH-AE, the structure model of was first predicted by AlphaFold^54^, and the model with high confidence (including one transmembrane helix) was manually fitted into the corresponding 4.4 Å focused refinement maps density map. Because the sidechains of NdhT cannot be identified under the current resolution, we use sequence poly-Ala in the NdhT model temporarily. All the subunits in the NDH-PSI model were rebuilt and adjusted manually using Coot^70^. After model building, real-space refinements were performed using Phenix-v1.17.1^73^, and the geometric restraints of the cofactors and the Chl– ligand relationships were supplied during the refinement. Automatic real-space refinements followed by manual correction in Coot were carried out interactively. PSI-1, PSI-2, NDH-AE and NDH-MBL were firstly refined against the masked maps, and then docked into the overall map and further refined. The geometries of the final structures were assessed using MolProbity^74^. A summary of the statistics for data collection and structure refinement of NDH-PSI supercomplexes from *Arabidopsis* is provided in Table S1. High-resolution images for publication were prepared using Chimera and PyMOL (Molecular Graphics System, LLC).

